# A high-fat diet induces a microbiota-dependent increase in stem cell activity in the *Drosophila* intestine

**DOI:** 10.1101/604744

**Authors:** Jakob von Frieling, Muhammed Naeem Faisal, Femke Sporn, Roxana Pfefferkorn, Stella Solveig Nolte, Felix Sommer, Philip Rosenstiel, Thomas Roeder

**Affiliations:** Zoological Institute, Department of Molecular Physiology, Kiel University, Kiel, Germany; IKMB, UKSH, Kiel University, Kiel, Germany; University Faisalabad, Pakistan; University Aarhus, Denmark; German Center for Lung Research, Airway Research Center North, Kiel, Germany

## Abstract

Over-consumption of high-fat diets (HFDs) is associated with several pathologies. Although the intestine is the organ that comes into direct contact with all diet components, the impact of HFD has mostly been studied in organs that are linked to obesity and obesity related disorders. We used *Drosophila* as a simple model to disentangle the effects of a HFD on the intestinal structure and physiology from the plethora of other effects caused by this nutritional intervention. Here, we show that a HFD triggers activation of intestinal stem cells in the *Drosophila* midgut. This stem cell activation was transient and preceded by induction of JNK signaling in enterocytes. JNK (basket) within enterocytes was essential for activation of stem cells in response to a HFD. Moreover, this nutritional intervention leads to upregulation of the cytokine *upd3* in enterocytes, a reaction that is known to trigger STAT signaling in intestinal stem cells followed by their activation. We also showed that the indigenous microbiota was essential for HFD-induced stem cell activation because this response was blunted in germ-free flies lacking a microbiota. Moreover, chronic HFD feeding of flies markedly altered both the microbiota composition and bacterial load. Although HFD-induced stem cell activity was transient, long-lasting changes to the cellular composition, including a substantial increase in the number of enteroendocrine cells, were observed. Taken together, a HFD enhances stem cell activity in the *Drosophila* gut and this effect is completely reliant on the indigenous microbiota and also dependent on JNK signaling within intestinal enterocytes.

**Author summary:** High-fat diets have been associated with a plethora of morbidities. The major research focus has been on its effects on obesity related disorders, mostly omitting the intestine, although it is the organ that makes the first contact with all diet components. Here, we aimed to understand the direct effects of HFD on the intestine itself. Using *Drosophila* as a model, we showed that HFD induced a transient activation of intestinal stem cells. This response completely depended on JNK signaling in enterocytes. Mechanistically, we found that HFD induces JNK signaling in enterocytes, which triggers production of the cytokine *upd3*. This, in turn activates STAT signaling in intestinal stem cells, leading to their activation. Surprisingly, we found that a HFD triggered induced stem cell activation critically depends on the indigenous microbiota, as in germ free flies this reaction was completely abolished. Moreover, HFD changed both, composition and abundance of the microbiota. As fecal transfer experiments failed to recapitulate the HFD phenotype, we assume that the increased bacterial load is the major cause for the HFD triggered stem cell activation in the intestine.

## Introduction

High caloric intake and especially high-fat diets (HFDs) are major causes of the epidemic increases in obesity-associated diseases [1]. In addition to metabolically relevant organs, the intestines are particularly susceptible to the effects of HFDs because they are in direct contact with constituents of the diet. Consequently, nutritional interventions directly impact intestinal structure and functionality [2]. Diet-dependent plasticity in the size of the intestines has been reported [3]. Specifically, intestinal size decreases in response to dietary restriction [4], but increases upon re-feeding after a period of starvation [5]. Structural changes in response to HFDs are observed at different levels, ranging from subcellular structures in enterocytes [6] to the cellular composition of the intestinal epithelium [7]. The effects of HFDs on the intestinal structure involve alteration of the activity of intestinal stem cells (ISCs). In mammals, HFDs directly enhance the activity of ISCs, leading to increased villi lengths in the small intestines via a mechanism involving ß-catenin signaling [8]. This HFD-induced activation of ISCs appears to be directly caused by the lipid content of food [3, 9]. A recent study showed that food with high lipid contents induces very robust PPAR-δ activation in ISCs and thereby increases the number of mitotically active cells in the intestines [9, 10]. Moreover, this response increases the tumorigenicity of intestinal progenitors [2, 9]. This observation correlates with epidemiological studies showing that different types of diets affect the risk of developing intestinal cancers [11, 12]. Specifically, HFDs increase the prevalence of colon cancers [13, 14]. In this context, deregulated stem cell activities appear to be directly associated with alterations in intestinal JAK/STAT signaling [15].

Although the major effects of HFDs on the stemness and tumorigenicity of ISCs seem to be directly mediated by exposure of intestinal epithelial cells to fat components [9], secondary effects are also highly relevant to induction of the complex phenotype that results from chronic consumption of a HFD. One important link between HFDs and disease development is the intestinal microbiota [16–18]. High-fat dietary supplementation alters the abundance, composition, and physiological performance of the microbiota [19–21]. In flies, HFDs increase the microbial abundance in the intestines [22]. Associations between an altered or dysbiotic microbiota composition and metabolic diseases have been repeatedly reported [23, 24]. Furthermore, microbiota transfer experiments revealed that altered microbiota compositions play a causative role in the development of metabolic disorders [25, 26]. In *Drosophila*, a link between a dysbiotic microbiota and the activity of ISCs was reported [27]. In this context, age-associated increases in the abundance of a particular microbial colonizer and the infection of specific pathogens trigger induction of stem cell activities [28, 29].

Despite this large body of informative studies, knowledge of the mechanisms by which HFDs regulate the activity of ISCs is not comprehensive. Here, we used the fruit fly *Drosophila* as a model and showed that a HFD induces a transient increase in stem cell activity via JNK-dependent activation of cytokine expression in enterocytes. This effect is dependent on the microbiota. Thus, we speculate that HFDs elicit physiological effects not only directly via activating stem cells through exposure to different fat components, but also indirectly via altering the microbiota, especially the bacterial abundance in the intestines.

## Results

We used the fruit fly *Drosophila melanogaster* as a model to study the effects of a HFD on the structural and physiological characteristics of the intestines. Furthermore, we analyzed the contribution of the microbiota to these alterations. HFD feeding triggered a burst of stem cell activity in the midgut (Fig. 1). We used flies that expressed GFP under the control of the escargot (esg) driver, meaning expression was restricted to ISCs and their direct descendants, i.e., enteroblasts. This analysis revealed that in comparison with flies fed a control diet (CD) (Fig. 1A), short durations of HFD feeding induced substantial expansion of these cell types (Fig. 1B). Quantitative analysis of these cells was performed using flies that expressed luciferase in the same spatial pattern (Fig. 1C). Luminescence increased upon very short durations of HFD feeding, but it decreased over time (Fig. 1C). A similar increase in cell number upon HFD feeding was observed when only enteroblasts were analyzed using flies that expressed GFP under the control of the su(H)GBE-Gal4 driver (Fig. 1D–F). After 3 days, more enteroblasts were detected in flies fed a HFD (Fig. 1E) than in flies fed a CD (Fig. 1D). Similar results were obtained by quantitative analysis of luciferase expression (Fig. 1F). As observed with esg-specific signals (i.e., ISCs plus enteroblasts), luminescence remained elevated after 7 days of HFD feeding. The effects of different diets on ISC activity were directly evaluated by counting the number of phospho-H3 (pH3)^+^ cells in the gut. The number of pH3^+^ cells was increased after 3 days of HFD feeding (Fig. 1H) in comparison with flies fed a CD (Fig. 1G), which was further supported by a quantitative analysis of the numbers of pH3^+^ cells (Fig. 1I). To evaluate the effects of a HFD on the cellular composition of the intestines, we counted the number of enteroendocrine cells, which are direct descendants of enteroblasts, similar to enterocytes (Fig. 1K). A HFD increased the number of enteroendocrine cells in a time-dependent manner (Fig. 1L). This increase started at day 5, peaked at day 7, and was sustained for more than 2 weeks in comparison with flies fed a CD.

**Figure 1:**
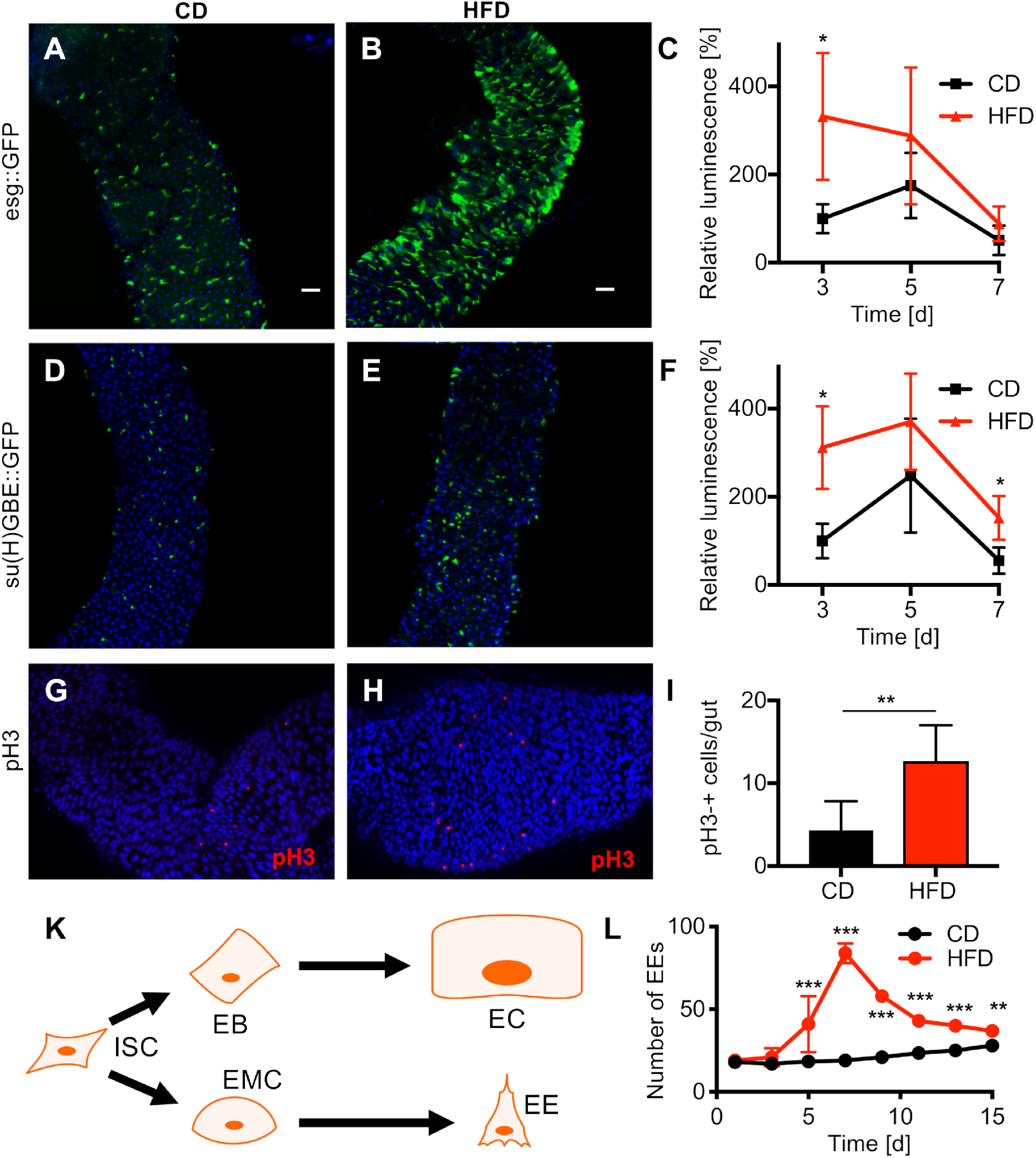
A HFD affects the cellular composition in the *Drosophila* intestine. (A–B) Intestines (anterior midgut R2 region) of flies with labeled ISCs and enteroblasts (*esg-Gal4::UAS-GFP*) and fed a CD (A) or a HFD for 3 days (B). (C) Luciferase signals of esg^+^ cells ectopically expressing luciferase (*esg-Gal4::UAS-luciferase*). Intestines were dissected from flies fed a CD or HFD for 3, 5, and 7 days (n = 9–14). (D–E) Pattern of GFP expressed under the control of the enteroblast-specific *su(H)GBE* driver (*su(H)GBE-Gal4::UAS-mCD8-GFP*) in intestines of flies fed a CD (D) or HFD (E). (F) Quantification of luciferase signals in su(H) GBE^+^ cells of flies fed a CD or HFD. (G–H) Representative images of anti-pH3 staining in the R4 region of the intestines in female flies fed a CD (G) or HFD (H). (I) Quantification of the number of pH3^+^ cells in whole intestines (n = 6). (K) Scheme of the generation of various cell types in the *Drosophila* midgut. ISC = intestinal stem cell, EB = enteroblast, EC = enterocyte, EMC = enteroendocrine mother cell, and EE = enteroendocrine cell. (L) Number of enteroendocrine (EE) cells in the R2 region of the intestines over 15 days in flies fed a HFD or CD (n = 10). DAPI was used as a counterstain in all images (scale bar = 50 μm). CD = control diet, HFD = high-fat diet. *p<0.05, **p<0.01, ***p<0.001.

JNK signaling is a major stress-sensitive signaling pathway in the intestines. Therefore, we tested if a HFD affects activation of JNK signaling (Fig. 2). In comparison with flies fed a CD (Fig. 2A), the level of phosphorylated JNK (pJNK), which corresponds to activated JNK, was increased in flies fed a HFD (Fig. 2B). Furthermore, using a JNK reporter line [30] (4xTRE-dsRed), we showed that the level of fluorescence in the anterior midgut was significantly higher in flies fed a HFD than in flies fed a CD (Fig. 2C). The cytokine upd3, which is directly linked to stem cell activation, is a major target of JNK signaling in enterocytes [31]. Thus, we investigated the effects of a HFD on expression of upd3. Using a reporter line (upd3-Gal4::UAS-GFP), we showed that, in comparison with flies fed a CD (Fig. 2D), expression of *upd3* in enterocytes was increased in flies fed a HFD (Fig. 2E). qRT-PCR revealed that the transcript level of *upd3* was significantly increased in flies fed a HFD (Fig. 2F). To determine whether HFD-induced JNK activation is causally linked with upd3 activation, we investigated if inhibition of JNK signaling via ectopic overexpression of a dominant-negative basket allele (*bsk^DN^*) in enterocytes affected *upd3* expression. HFD feeding did not increase *upd3* expression in flies expressing *bsk^DN^* in enterocytes (Fig. 2F), implying that activation of JNK upon HFD feeding is responsible for the increase in local *upd3* production and consequently stem cell activation. It is well established that upd3 produced by enterocytes induces proliferation of ISCs via activation of JAK/STAT signaling. Therefore, we used a STAT reporter line, in which GFP expression indicates JAK/STAT pathway activation [32]. As expected, GFP was detected in cells with a stem cell-like appearance scattered throughout the intestines. Fluorescence was relatively weak in flies fed a CD (Fig. 2G), but was far stronger in flies fed a HFD (Fig. 2H). Semi-quantitative analysis revealed that the level of fluorescence was significantly increased in flies fed a HFD (Fig. 2I).

**Figure 2.**
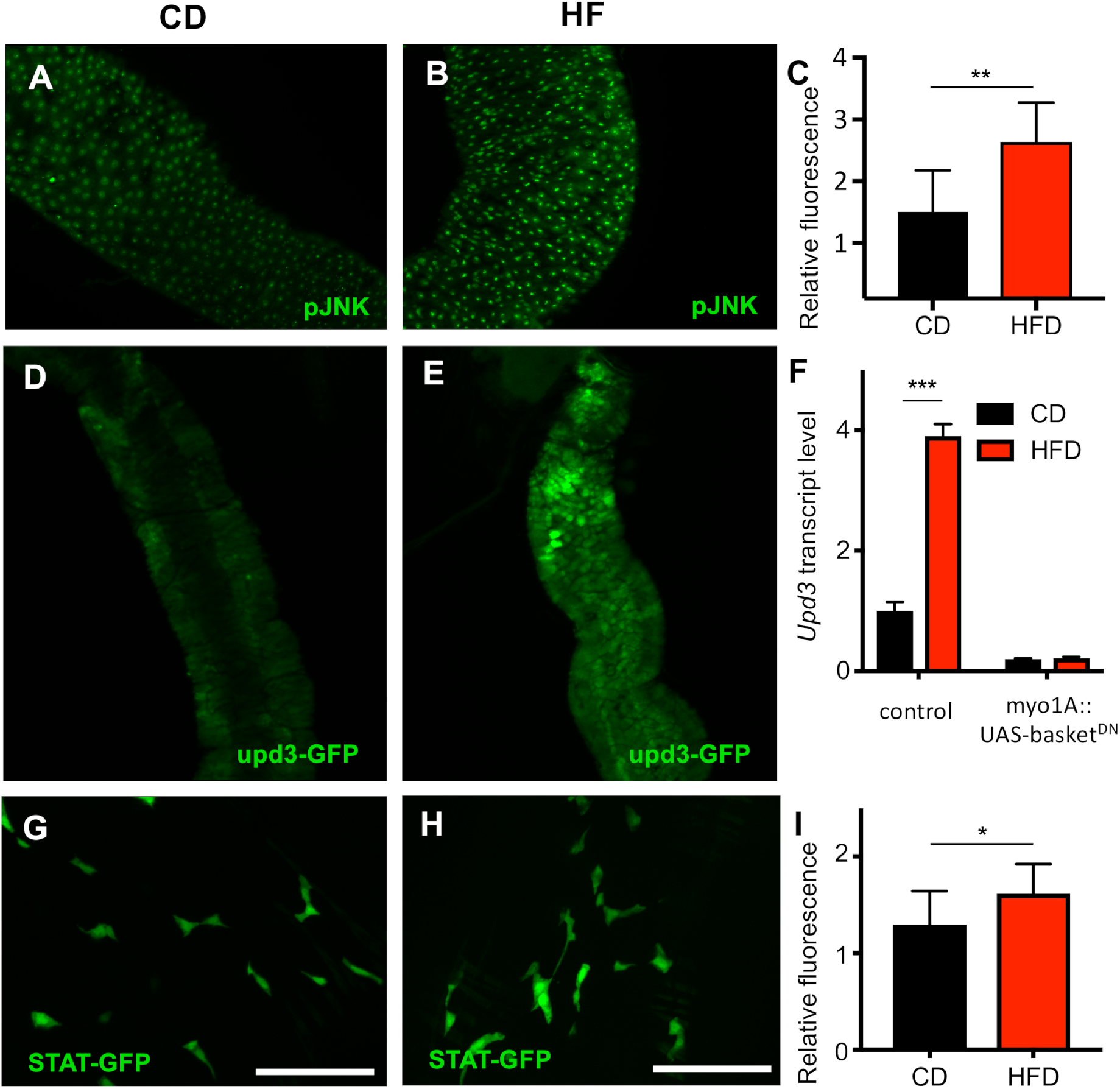
A HFD induces *upd3* expression via JNK signaling. (A–B) Staining with an anti-pJNK antibody in intestines of flies fed a CD (A) or a HFD for 3 days (B). (C) Fluorescence in the anterior midgut of a JNK reporter line (4XTRE-DsRed) fed a CD or HFD (n = 5). (D–E) An *upd3-GFP in vivo* reporter was used to monitor upd3 expression in the intestines of flies fed a CD (D) or a HFD for 3 days (E). Representative images of the anterior midgut R2 region of the intestines are shown. (F) qRT-PCR analysis of *upd3* expression in intestines of female flies that expressed a dominant-negative form of basket in enterocytes (*myo1A-Gal4::UAS-basket^DN^*) or the control (*w^1118^::UAS-basket^DN^*) and fed a CD or a HFD for 3 days (n = 5). (G–H) Images of intestines isolated from a *STAT-GFP in vivo* reporter strain fed a CD (G) or HFD (H) (scale bar = 50μm). (I) Quantification of fluorescence in the *STAT-GFP* reporter strain fed a CD or HFD for 3 days (n = 11). CD = control diet, HFD = high-fat diet. *p<0.05, **p<0.01, ***p<0.001.

Dietary interventions affect the indigenous microbiota. Therefore, we compared germ-free (GF) flies with those that had been reconstituted with a natural microbiota (Fig. 3). The latter flies had been used in all previously described experiments. As mentioned earlier, a HFD induced stem cell proliferation, which could be visualized by labeling ISCs and enteroblasts with GFP using *esg*-Gal4::UAS-GFP. In flies reconstituted with a natural microbiota, there were fewer *esg*^+^ cells in flies fed a CD (Fig. 3A) than in flies fed a HFD (Fig. 3B). By contrast, among GF flies, the number of *esg*^+^ cells was low in flies fed a CD (Fig. 3C) or HFD (Fig. 3D). Similar results were obtained by quantitative analysis of the number of *esg*^+^ cells (Fig. 3E). This lack of induction in GF flies was also observed using reporter strains that allowed visualization of *upd3* expression (Fig. 3F–J), indicating that GF flies lack the signal necessary to induce proliferation of ISCs upon HFD feeding. Whereas the upd3 signal was low in flies reconstituted with a native microbiota and fed a CD (Fig. 3F), it was sustainably increased upon HFD feeding (Fig. 3G). By contrast, the upd3 signal was very low in GF flies fed a CD (Fig. 3H) or HFD (Fig. 3I). qRT-PCR revealed that the transcript level of *upd3* was significantly lower in GF flies fed a HFD or CD than in flies reconstituted with a functional microbiota (Fig. 3J).

**Figure 3.**
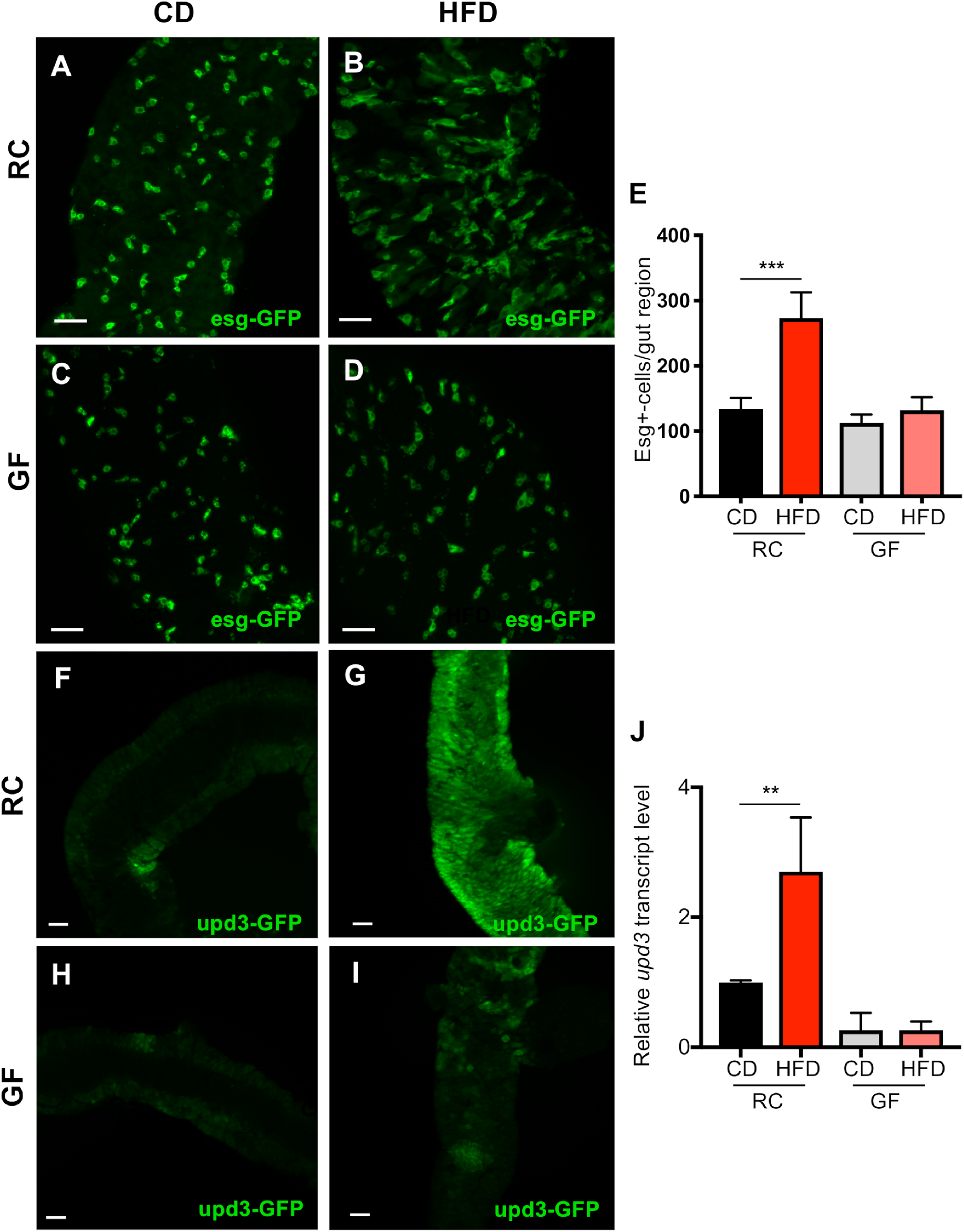
HFD-induced cell proliferation and upd3 expression are dependent on the intestinal microbiota. (A–D) The *esg-Gal4::UAS-GFP* strain, in which ISCs and enteroblasts were labeled, was examined. Flies reconstituted with a normal microbiota were fed a CD (A) or HFD (B). GF flies were fed a CD (C) or HFD (D). Representative images of the anterior midgut R2 region are shown (scale bar = 50 μm). (E) Number of *esg^+^* cells in the R2 region of the intestines (n = 10). (F–I) An *upd3-GFP* reporter was used to examine microbiota-associated modulation of upd3 expression. Flies reconstituted with a normal microbiota were fed a CD (F) or HFD (G). GF flies were fed a CD (H) or HFD (I). Representative images of the R2 region are shown (scale bar = 50 μm). (J) qRT-PCR analysis of *upd3* expression in GF and recolonized flies fed a CD or HFD (n = 5). CD = control diet, HFD = high-fat diet, GF = germ-free, RC = recolonized. **p<0.01, ***p<0.001.

To elucidate the effects of a HFD on the microbiota, we compared the microbiota composition between flies fed a HFD and those fed a CD. A HFD significantly altered the composition of the microbial community in comparison with a CD, as illustrated by the separation in beta diversity by principal coordinate analysis based on unweighted UNIFRAC (Fig. 4A). Linear discriminant analysis effect size (LEfSe) revealed that the orders Enterobacteriales and Caulobacterales were significantly enriched upon HFD feeding. On the other hand, flies fed a CD exhibited enrichment of species belonging to the family Lactobacillaceae, especially from the genus *Pediococcus* (Fig. 4B). To determine whether these alterations in the microbial community were sufficient to induce stem cell activity, we performed a fecal transplantation assay (Fig. 4C–G). Specifically, we transferred the microbiota of flies fed a CD or HFD into GF flies expressing GFP under the control of the esg driver. Transfer of the microbiota from flies fed a CD did not affect the number of esg^+^ cells after 3 days (Fig. 4C) or 5 days (Fig. 4D). In addition, transfer of the microbiota from flies fed a HFD did not increase the number of these cells after 3 days (Fig. 4E) or 5 days (Fig. 4F). Quantitative analysis of esg^+^ cells revealed that neither transfer affected stem cell activity in the intestines (Fig. 4G), indicating that alteration of the microbiota composition does not underlie the increase in stem cell activity observed upon HFD feeding.

**Figure 4.**
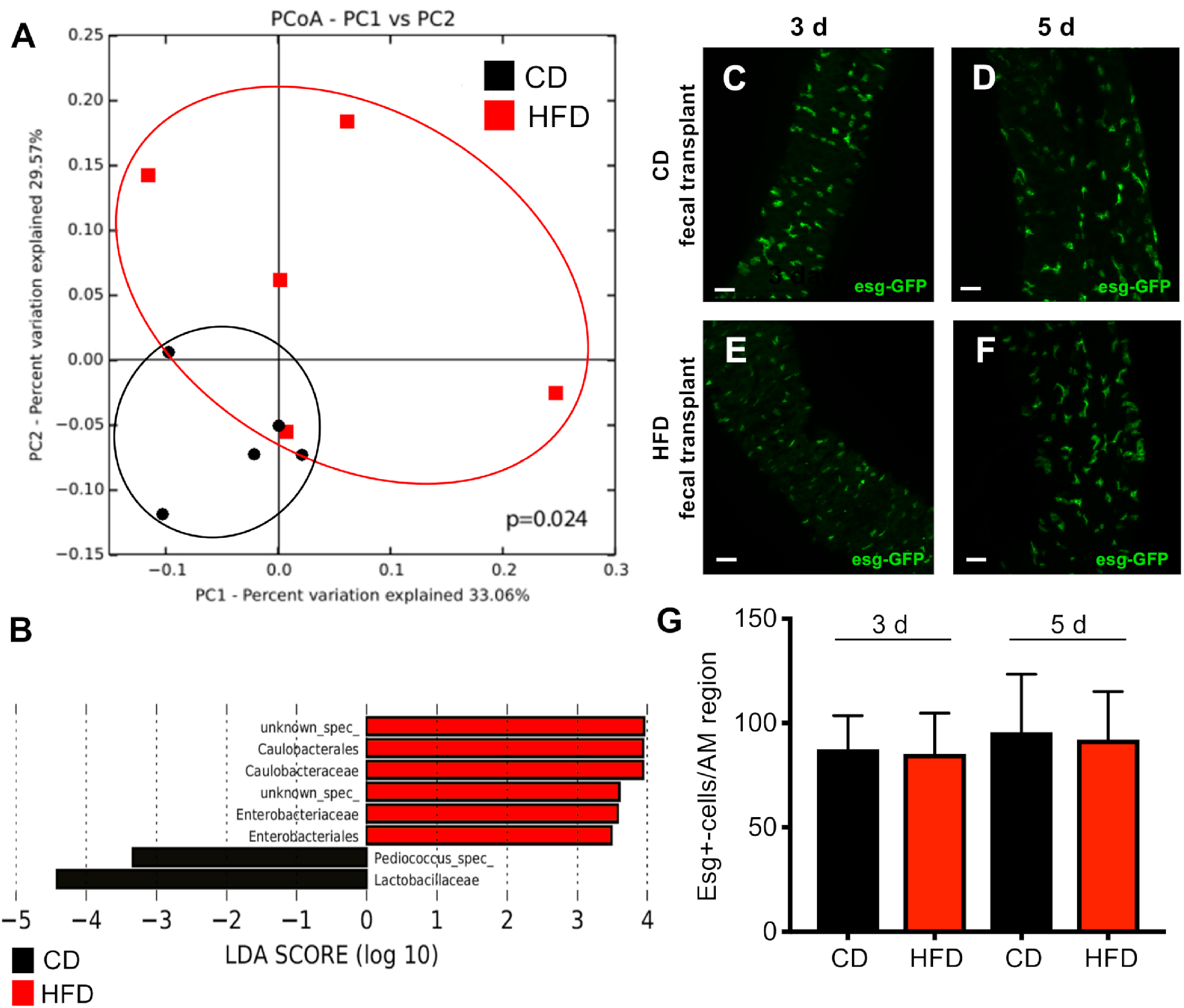
A HFD alters the microbial composition in w^1118^ flies. (A) Principal coordinate analysis showing significant alterations in the intestinal microbiota triggered by a CD or HFD. Each data point corresponds to one biological replicate. (B) Linear discriminant analysis effect size (LEfSe) to determine differentially enriched bacteria in the intestinal microbiota of flies fed a CD or HFD. (C–F) Fecal transplantation into the *esg-Gal4::UAS-GFP* reporter strain was performed to examine the effect of diet-induced changes in the microbial composition on intestinal cell proliferation. Flies were reconstituted with a microbiota derived from flies fed a CD (C–D) or HFD (E–F). Samples were analyzed after 3 days (C and E) or 5 days (D and F) (scale bar = 50 μm). (G) Quantitative analysis of the data presented in (C–F) (n = 8–12). CD = control diet, HFD = high-fat diet.

As well as altering the microbiota composition, a HFD changed the bacterial load (Fig. 5). The bacterial load in the intestines was ~2-fold higher in flies fed a HFD than in flies fed a CD (Fig. 5A). Furthermore, a HFD induced mild constipation and consequently the number of fecal spots deposited over 24 h was substantially reduced by ~60–70% (Fig. 5B). In addition, the number of deposited fecal spots was almost 50% lower in GF flies than in flies reconstituted with a functional microbiota. A HFD significantly reduced fecal spot production both in flies reconstituted with a functional microbiota and GF flies. The gut transit time was shortest in flies reconstituted with a functional microbiota and fed a CD, but was increased in flies fed a HFD (Fig. 5C). This effect was more pronounced in GF flies. The intestinal diameter was larger in flies fed a HFD than in flies fed a CD (Fig. 5D-F), which may reflect the increased bacterial mass in the intestines.

**Figure 5.**
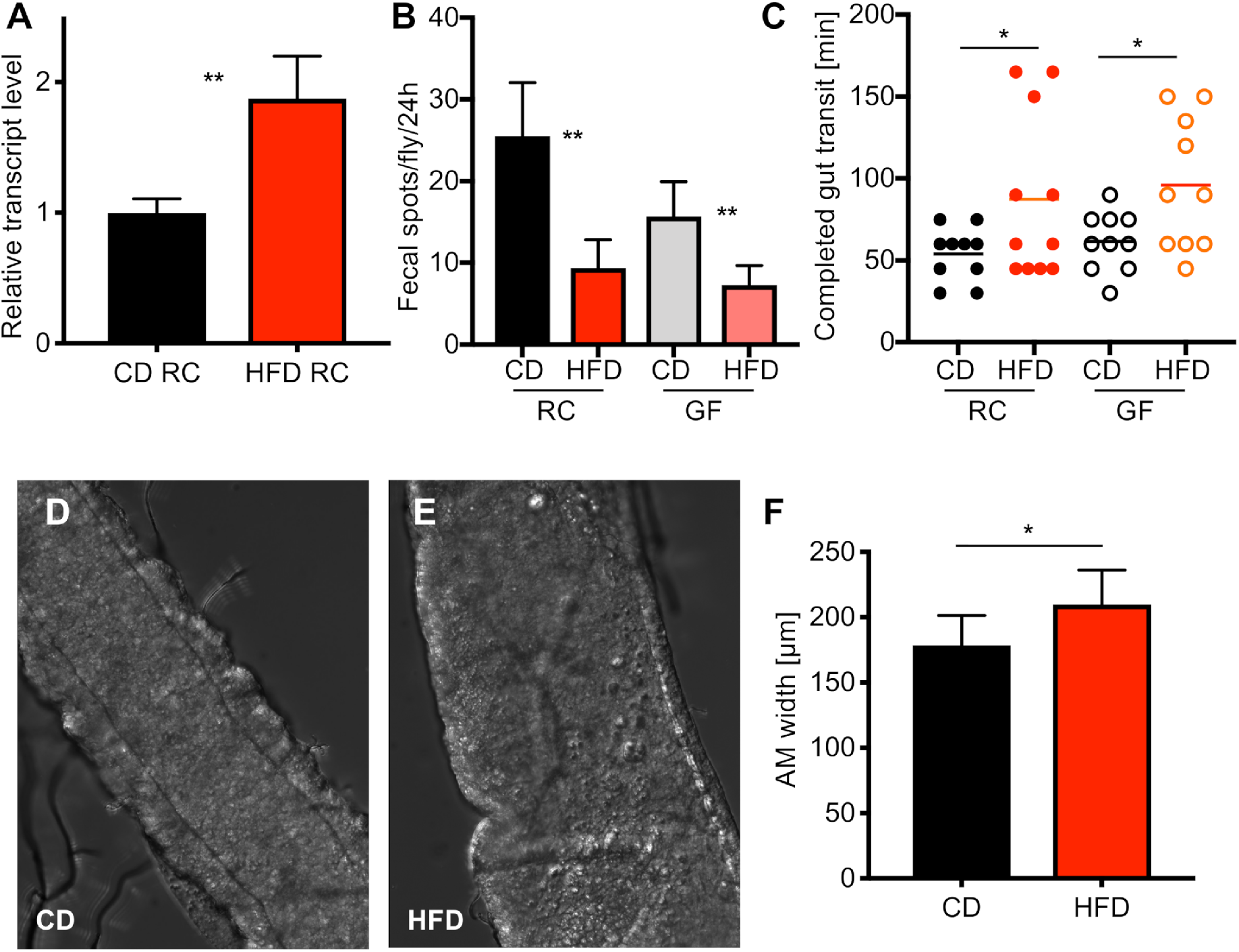
A HFD affects egestion and microbial abundance. (A) The intestinal bacterial load was analyzed by qRT-PCR using 16S universal primers. Intestines were dissected from flies fed a CD or HFD for 3 days (n = 5). (B) Analysis of fecal spot production over 24 h in flies with a reconstituted microbiota and GF flies fed a CD or HFD (n = 7–10). (C) Gut transit time, defined as the time from food ingestion to egestion, was assessed by feeding flies blue food and determining the time until excretion of blue feces (n = 10). (D–E) Representative images of the R2 region of flies fed a CD (D) or a HFD for 3 days (E). (F) Quantification of intestinal thickness (n = 11). CD = control diet, HFD = high-fat diet, GF = germ-free, RC = recolonized. *p<0.05, **p<0.01.

## Discussion

High-fat or lipid-rich diets are thought to be responsible for the development of a plethora of metabolic disorders, including the epidemic increases in obesity and type 2 diabetes. The current study focused on the effects of a HFD on the intestines, which is the first organ exposed to this dietary intervention. We investigated the effects of a HFD on intestinal structure and physiology using *D. melanogaster* as a simple model. We revealed that a HFD induced a substantial increase in stem cell activity in the intestines, leading to altered cell numbers and changes in the cellular composition of this organ. This induction of stem cell activation was completely reliant on a functional intestinal microbiota. A HFD did not increase stem cell activity in GF flies. Moreover, this effect was also dependent on JNK signaling in enterocytes. We assume that JNK signaling transduces this microbiota-dependent signal in enterocytes, leading to production and release of the cytokine upd3. This, in turn, activates JAK/STAT signaling in ISCs and is thus essential for induction of proliferation in these cells. Taken together, activation of JNK signaling in enterocytes upon HFD feeding is completely reliant on the intestinal microbiota. The effects of HFDs on the structural and functional characteristics of the intestines have been reported in several models. Interestingly, different mechanisms appear to underlie the effects of HFDs on stem cell proliferation. In *Drosophila*, diets containing very high cholesterol concentrations significantly increase the abundance of enteroendocrine cells. This is due to a direct interaction with Notch signaling, which leads to preferential production of enteroendocrine cells [7]. Moreover, reduction of the lipid content decreases the proliferation of enteroendocrine tumors. A HFD induces comparable effects in mice, which are mainly characterized by increased cell proliferation in the intestines [33–35]. Beyaz and colleagues [9, 10] reported that a HFD induces stemness of ISCs, which is accompanied by decoupling of stem cell activity from its niche. This effect is opposite to the augmentation of stem cell function observed in response to dietary restriction in the intestines, which is dependent on increased integration into the stem cell niche rather than decoupling [4]. Comparable with the effects of cholesterol in the intestines of *Drosophila*, the effects of a HFD can be traced back to the direct interactions of particular fatty acids with ISCs [10].

These previously reported effects of HFDs on stemness are due to direct interactions between specific compounds in these diets (cholesterol or fatty acids) and intestinal cells. This is in contrast with the mechanism by which a HFD promotes stemness in our system, which is indirect because it requires the microbiota. HFDs are associated with a dysbiotic microbiota, which has been causally linked with various pathogenic states. The microbiota is also dysbiotic during aging [27, 28]. Most previous studies focused on the microbiota composition and reported that a shift toward specific bacterial colonizers correlates with the potential for disease development in the host. This appears to be irrelevant in *Drosophila* because fecal transfer experiments failed to recapitulate the enhanced stemness phenotype. Instead, an increased abundance of bacterial colonizers appears to be more relevant. Two factors likely contribute to the increased microbial abundance in response to a HFD: 1) the high-energy content, which facilitates bacterial growth within the intestines, and 2) the increased gut transit time, which reduces fecal output and thus loss of bacteria. The gut transit time was increased and fecal output was reduced in flies fed a HFD, and these changes might underlie the HFD-induced increase in microbial abundance. An increased gut transit time upon HFD feeding is also observed in mice and humans [36–38], indicating that it is a general response to this nutritional intervention and that a HFD is usually associated with an increased bacterial abundance. One consequence of the increased microbial abundance is elevated pressure within the intestinal lumen, leading to mechanical stress on the intestinal structure. This provides another scenario to explain how a HFD increases the stemness of ISCs; specifically, increased pressure induces an increase in stem cell activity in the intestines [39]. The piezo channel in precursors of enteroendocrine cells responds to mechanical stress, and this leads to production of enteroendocrine cells, which is consistent with the results of this study. We also showed that JNK signaling in enterocytes is essential for a suitable cellular response to such stress [40, 41]. This sentinel function of enterocytes is highly relevant because these cells are confronted with a plethora of stress signals and must signal about their state to enable their replacement if necessary [42, 43].

In addition to the effects of a HFD on stem cell activity in the intestines, we also found that this dietary intervention induces long-lasting modifications of the intestinal hormonal architecture. A HFD increased the number of enteroendocrine cells for a considerable duration. This observation is consistent with previous reports that a HFD has multiple effects on intestinal function and homeostasis [44] and changes the expression and release of gut hormones [45]. These hormones play a central role in metabolic control and regulate various aspects of intestinal function [46]. We cannot rule out the possibility that feeding of a HFD for even a relatively short duration has long-lasting effects on intestinal homeostasis. In mammals, the substantial and long-lasting effects of short episodes of HFD feeding (e.g., only 3 days) attenuate major lipid-sensing systems in the gut [47]. HFDs usually reduce appetite via hormonal circuitry [48] to effectively prevent overnutrition [49]. Whereas almost nothing is known about the effects of HFDs on release of gut hormones in *Drosophila*, a plethora of studies have reported such effects in various mammals [50, 51]. Chronic exposure to HFDs increases release of the major gut hormone cholecystokinin in rats [52]. Similar effects have been reported for other gut hormones such as glucagon-like peptide 1 [53]; however, the underlying mechanism remains to be elucidated. Few interventions have been shown to change the number of enteroendocrine cells in the intestines [54]. Such a change would not only modify hormonal and metabolic homeostasis, but may also alter the stem cell niche because enteroendocrine cells are highly relevant for maintenance of this important niche [55].

Taken together, our results clearly show that a HFD elicits major effects on intestinal structure and function even in the simple model organism *D. melanogaster*. These effects include transient activation of stem cell activity and long-lasting changes to the cellular architecture in the intestines. Moreover, these effects are completely dependent on the microbiota. The stress-sensing JNK signaling pathway appears to be of central importance in the mechanism underlying these effects.

## Material and Methods

### Fly food and husbandry

All flies were raised in vials on sterile standard cornmeal medium containing 5% inactivated yeast (BD Bacto™ yeast extract), 8.6% cornmeal (Mühle Schlingemann, Waltrop, Germany), 5% glucose (Roth, Karlsruhe, Germany), and 1% agar-agar (Roth) in an incubator at 20°C and 65% humidity. At 3–5 days after hatching, adult flies were transferred to fresh sterile standard cornmeal medium or high-fat medium. High-fat medium was exactly the same as standard cornmeal medium except that it contained 20% (w/v) food-grade palm fat (Palmin^®^). The following fly strains were used in this study: *w^1118^* (Bloomington Stock Center), *esg-GFP* (gift from N. Perrimon, Harvard University, USA), amon-Gal4 (gift from C. Wegener, Würzburg, Germany), *UAS-Luc* (gift from M. Markstein, University of Massachusetts, USA), *su(H)GBE-Gal4* (gift from S. Hou, Frederick, USA) *UAS-basket^DN^* (Bloomington Stock Center 44801), *10XSTAT::GFP* (gift from E. Bach, New York, USA), *20xUAS-IVS-mCD8::GFP* (Bloomington Stock Center 32194), and 4XTRE-DsRed (Bloomington Stock Center 59011).

### Axenic flies

Flies were allowed to lay eggs on apple juice agar plates containing 2% agar-agar (Roth 5210.2) and 50% apple juice (Rewe Bio apple juice) for about 12 h at 25°C to prevent the presence of larvae. Eggs were collected by rinsing the apple juice agar plates with deionized water and transferred to net baskets. Thereafter, the eggs were bleached for 2 min with 6% sodium hypochlorite (Roth, Karlsruhe, Germany) and then washed with 70% ethanol (Roth) and double autoclaved water under sterile conditions. Bleached embryos were placed on sterile standard cornmeal medium. The lack of bacteria in emerged adults was tested by PCR using universal primers targeting bacterial 16S rDNA.

### Reconstitution of bacteria

A mixture of the following bacterial species, which were cultured as previously described [56], was used to reconstitute the natural microbiota: *Acetobacter pomorum* (OD_600_ = 0.7), *Lactobacillus brevis^EW^* (OD_600_ = 8), *Lactobacilles plantarum^WJL^* (OD_600_ = 6), *Enterococcus faecalis* (OD_600_ = 0.8), and *Commensalibacter intestini^A911T^* (OD_600_ = 1.5) (kindly provided by Carlos Ribeiro, Lisbon, Portugal). To prepare the stock solution for recolonization, the following volumes of each liquid culture were combined in a 15 ml falcon tube: 2 ml of *A. pomorum*, 0.02 ml of *L. brevis^EW^*, 0.25 ml of *L. plantarum^WJL^*, 2 ml of *E. faecalis*, and 1 ml of *C. intestini^A911T^*. The bacterial mixture was centrifuged three times at 3000 rpm for 15 min and repeatedly resuspended in sterile phosphate-buffered saline (PBS). Finally, the mixture was centrifuged at 3000 rpm for 15 min and resuspended in 25% glycerol prepared in sterile PBS. Aliquots of 500 μl were stored at −20°C. A volume of 50 μl of the bacterial mixture was added to the surface of control or high-fat media and allowed to settle for 1 h at room temperature under sterile conditions. Thereafter, 3–5-day-old GF flies were transferred to the corresponding media.

### Fecal transplantation

Reconstituted esg-GFP flies aged 3–5 days were cultured on high-fat or control media for 7 days. Subsequently, five flies were homogenized in 300 μl of MRS medium (BD Difco™, Thermo Scientific, Braunschweig, Germany) using a Bead Ruptor 24 (BioLab Products, Bebensee, Germany). Fly debris was removed by centrifugation, the supernatant was mixed with 5% sucrose, and 200 μl of the mixture was applied to sterile filter paper. GF *esg*-GFP flies were allowed to feed on the filter paper for 24 h and then transferred to control media for 5 or 7 days. GFP^+^ cells in the anterior midgut region were counted.

### Body fat quantification

Total body triacylglycerols in flies were measured using a coupled colorimetric assay as described previously [18, 57]. Groups of five females were weighed and homogenized in 1 ml of 0.05% Tween-20 using a Bead Ruptor 24 (BioLab Products, Bebensee, Germany). Homogenates were heat-inactivated at 70°C for 5 min, centrifuged, and incubated with triglyceride solution (Thermo Fisher Scientific, Braunschweig, Germany) at 37°C for 30 min. A standard curve was prepared using glyceryl trioleate. Absorbance at 562 nm was measured.

### Immunohistochemistry

Intestines were dissected in PBS, fixed in 4% paraformaldehyde for 1 h at room temperature, washed three times with PBST (PBS containing 0.1% Triton X-100), and blocked in blocking buffer (PBST containing 5% normal goat serum) for 1 h at room temperature. Thereafter, intestines were incubated with a primary antibody diluted in blocking buffer overnight at 4°C, washed three times, and then incubated with a secondary antibody diluted in PBST overnight at 4°C in darkness. Subsequently, intestines were washed three times with PBST and mounted on slides in Mowiol 40-88. Images were acquired using a fluorescence microscope equipped with Apotome (Carl Zeiss Image Axio Vision, Jena, Germany). The following antibodies were used: anti-prospero from mouse (1:50, Developmental Studies Hybridoma Bank, Iowa City, USA, MR1A), anti-GFP from mouse (1:300, Developmental Studies Hybridoma Bank, Iowa City, USA 8H11), anti-pJNK polyclonal from rabbit (Promega, Mannheim, Germany), Alexa Fluor 488-labeled goat anti-mouse IgG (1:300, Jackson ImmunoResearch, Dianova, Hamburg, Germany), and Alexa Fluor 555-conjugated goat anti-mouse IgG (1:300, Cell Signaling Technology, Frankfurt, Germany).

### Fluorescence-based quantification of the *in vivo* STAT-GFP reporter

Intestines of 10xSTAT::GFP flies were dissected after CD or HFD feeding for 3 days and immediately fixed in 4% paraformaldehyde and washed three times with 0.1% PBST. All images were stacked with a thickness of 2 μm and an exposure time of 60 ms. Fluorescence of all cells in the field of view was measured using ImageJ. Corrected total cell fluorescence (CTCF) was calculated as follows: CTCF = integrated density – (area of selected cell × mean fluorescence of background readings).

### Luciferase assay

The luciferase assay was performed as previously described [58] with minor modifications. The intestines of five adult flies per replicate were collected in 150 μl of Glo Lysis Buffer (Promega, Mannheim, Germany, #E2661) and homogenized using a bead mill homogenizer (BioLab Products, Bebensee, Germany) for 2 min at 3.25 m/s. The homogenate was transferred to a new reaction tube and stored at −20°C until further processing. For luciferase measurement, samples were thawed on ice and 50 μl was transferred to a white flat-bottom 96-well plate, with at least one empty well between treatments. Samples were mixed with the same amount of substrate provided by the One Glo Luciferase Assay System (Promega, Mannheim, Germany, #E6110) immediately before signal detection. Luciferase signals were detected using a Tecan plate reader (Tecan, Infinite M200 Pro, Männedorf, Switzerland). A defined control was included on each plate to normalize treatments across plates.

### Assessment of fecal output

A small piece of CD or HFD supplemented with Brilliant Blue FCF food dye (E133, Ruth, Bochum, Germany) was transferred to the bottom of a vial. A coverslip was placed in the middle of the vial to split it into two halves. Individual flies were trapped in one half, together with the piece of food, and the vial was sealed with a foam plug. Flies were incubated at 20°C for 24 h. Coverslips were scanned. The numbers of fecal spots was counted manually.

### Assessment of the intestinal transit time

Each well of a 24-well plate was loaded with a small piece of CD or HFD supplemented with Brilliant Blue FCF food dye. Individual flies were starved for 24 h, transferred to the wells, and monitored every 15 min for 3 h. The appearance of the first dyed fecal spot determined the time from food ingestion to egestion in each individual fly, which was referred to as the transit time.

### RNA extraction and qRT-PCR

Total RNA was extracted from the midgut of adult female flies that had been kept on standard cornmeal medium or high-fat medium. qRT-PCR was performed as described previously [59]. The following primers were used: *upd3* forward (5’-GAGAACACCTGCAATCTGAA-3’) and *upd3* reverse (5’-AGAGTCTTGGTGCTCACTGT-3’). The primers 8FM (5′-AGAGTTTGATCMTGGCTCAG-3′) and Bact515R (5′-TTACCGCGGCKGCTGGCAC-3′) were also used to quantify the bacterial load.

### 16S amplicon sequencing

DNA was isolated from intestines containing fecal material using a DNeasy Blood and Tissue Kit (Qiagen, Hilden, Germany) following the manufacturer’s protocol for purification of total DNA from animal tissues and for pretreatment of Gram-positive bacteria. Intestines of 10 individual flies were pooled per sample to generate sufficient material. Extracted DNA was eluted from the spin filter silica membrane with 100 μl of elution buffer and stored at −80°C.

16S profiling and MiSeq sequencing were performed as described previously [60, 61] with modifications. The V3-V4 region of the 16S gene was amplified using the dual barcoded primers 341F (GTGCCAGCMGCCGCGGTAA) and 806R (GGACTACHVGGGTWTCTAAT). Each primer contained additional sequences for a 12-base Golay barcode, an Illumina adaptor, and a linker sequence [62]. PCR was performed using Phusion Hot Start Flex 2× Master Mix (NEB, Frankfurt, Germany) in a GeneAmp PCR system 9700 (Applied Biosystems, Thermo Fisher Scientific, Frankfurt, Germany) and the following program: 98°C for 3 min, 30 cycles of 98°C for 20 s, 55°C for 30 s, and 72°C for 45 s, followed by 72°C for 10 min and then 4°C hold. PCR was checked by agarose gel electrophoresis. Normalization was performed using a SequalPrep Normalization Plate Kit (Thermo Fisher Scientific, Darmstadt, Germany) following the manufacturer’s instructions. Equal volumes of normalized amplicons were pooled and sequenced on an Illumina MiSeq (2 × 300 nt).

MiSeq sequence data were analyzed using MacQIIME v1.9.1 [63]. Briefly, all sequencing reads were trimmed to retain only nucleotides with a Phred quality score of at least 20, and then paired-end assembled and mapped onto the different samples using the barcode information. Rarefaction was performed at 34,000 reads per sample to normalize all samples against the minimum shared read count and to account for differential sequencing depth. Sequences were assigned to operational taxonomic units (OTUs) using UCLUST and the Greengenes reference database (gg_13_8 release) with 97% identity. Representative OTUs were picked and assigned to a taxonomy using UCLUST and the Greengenes database. Quality filtering was performed by removing chimeric sequences using ChimeraSlayer and by removing singletons and sequences that failed to align with PyNAST. The reference phylogenetic tree was constructed using FastTree 2. Relative abundance was calculated by dividing the number of reads for an OTU by the total number of sequences in the sample. Unweighted Unifrac beta diversity was calculated and visualized by generating principal coordinate plots. Differentially abundant taxa were assessed using the nonparametric t test. p values were adjusted for multiple testing using the FDR correction. LEfSe [64] was performed using an online tool available at http://huttenhower.sph.harvard.edu/galaxy. LDA denotes taxa based on their contribution to the overall observed differences between groups, i.e., taxa whose abundance was significantly higher in flies fed a HFD than in flies fed a CD.

### Statistical analysis

All statistical analyses were performed using Prism 6.0 (GraphPad Software, San Diego, USA). Lifespan data were analyzed by the log rank test (Mantel-Cox).

## Acknowledgments

We thank Britta Laubenstein and Christiane Sandberg for excellent technical help. In addition, we thank Michelle Markstein, Erika Bach, Norbert Perrimon, Stephen Hou, Chris Wegner, and the Bloomington Stock Center for providing flies. This work was supported by the German Research Foundation as part of CRC 1182 (subproject C2) and by the Cluster of Excellence ‘Inflammation at Interfaces’.

